# Motor unit discharge properties of the vastii muscles and their modulation with contraction level depend on the knee-joint angle

**DOI:** 10.1101/2024.11.29.625975

**Authors:** Tamara Valenčič, Sumiaki Maeo, Stefan Kluzek, Aleš Holobar, Jakob Škarabot, Jonathan P Folland

## Abstract

This study examined the effect of the knee-joint angle on motor unit (MU) discharge properties of the vastii muscles and their modulation with contraction level. Twelve young adults performed unilateral isometric knee-extension contractions during three experimental sessions at either 25, 55, and 85° of knee flexion (full extension: 0°) in a randomised order. Each session involved maximal voluntary contractions (MVCs) followed by submaximal trapezoidal and triangular contractions at different levels relative to maximal voluntary torque (MVT). High-density surface electromyograms were recorded from vastus lateralis and medialis muscles and, subsequently, decomposed to obtain discharge timings of individual MUs. MVT was the greatest, whereas MU discharge rate (DR) during MVCs and submaximal contraction levels (≥30% MVT) was the lowest at the intermediate joint angle (55°). The highest DR during MVCs and high-level contractions (70% MVT), however, was at the most flexed knee position (85°), which was due to a greater DR increase 50-70% MVT compared to 25° and 55°. The onset-offset DR hysteresis (ΔF), an estimate of persistent inward current contribution to motoneuron discharge, decreased with knee flexion and increased with contraction level, whereas the degree of motoneuron input-output nonlinearity (brace height) did not vary with joint angle but decreased with contraction level. At 85°, ΔF increased more and brace height decreased less with contraction level compared to 25° and 55°. These findings indicate that vastii MU DR and its modulation with contraction level vary with knee-joint angle, which could be partly explained by the modulation of motoneuron intrinsic electrical properties.

**NEW & NOTEWORTHY:** This study explored the relationship between motoneuron output to the vastii muscles at different knee-joint angles (quadriceps lengths) and isometric contraction levels. We showed that the motor unit discharge rate was lowest at the angle of the greatest absolute torque capacity, whereas the contraction-level-induced increases in discharge rate and motoneuron excitability were the greatest in the flexed position. These findings suggest that joint-angle-dependent adjustments in sensory feedback modulate motor control of the knee-extensor muscles.

## INTRODUCTION

Muscle torque output is determined by both the contractile properties and neural activation of the constituent motor units (MUs), including recruitment and discharge rate modulation. Whilst changes in joint angle, and thus muscle length, are known to influence the intrinsic contractile properties of a muscle (1), the effect on neural activation is less clear. Nonetheless, different joint angles are known to impact the strain on muscles (agonist and antagonist), tendons, ligaments, and other joint structures (2, 3), likely modulating the balance between excitatory and inhibitory sensory inputs to the spinal cord and supraspinal centres, and thus the neural strategies of muscle torque production (i.e., motoneuronal output) (4, 5).

In various upper- and lower-limb muscles, studies reported either similar motor unit discharge rate at all muscle lengths/joint angles (*tibialis anterior:* (6, 7), triceps brachii: (8)) higher at short (*biceps brachii:* (9); *hamstrings:* (10)) or higher at long (*triceps surae:* (11–13)) muscle lengths during submaximal isometric contractions performed at the same relative torque level (% maximal voluntary torque). Furthermore, when comparing the increases in MU discharge rate with contraction level at different joint angles/muscle lengths, either no difference (6, 7), larger increases at short (10) and larger increases at long (12) muscle lengths have been noted. Whilst some of these disparities may be attributed to differences in and limitations of the methodological approaches used across studies, it is also possible that joint-angle- and contraction-level-dependent adjustments in the neural control of torque production vary between muscles and joints.

The quadriceps femoris, the primary extensor of the knee joint, is the largest anatomical muscle group in the human body. Its force vector generates significant compression of both the tibiofemoral and patellofemoral joints (2, 14) and induces anterior tibial translation (15, 16), thereby straining the anterior cruciate ligament (ACL) (3). As the magnitude of the internal joint loads varies with contraction level and knee-joint angle (16), and modulates afferent feedback from the mechanosensitive muscle and joint receptors (17), it is plausible that the neural control of quadriceps torque production is more sensitive to changes in joint angle compared to smaller, weaker muscles controlling smaller joints (e.g., ankle and elbow). Supporting this hypothesis, several lines of evidence suggest voluntary activation of the quadriceps during maximal isometric knee-extension contractions varies with knee position. For example, global quadriceps activation assessed via surface electromyography (sEMG) or the interpolated twitch technique (18) during maximum voluntary isometric contractions has generally been found to decrease at extended compared to flexed knee positions (19–22). Similarly, quadriceps neuromuscular activity (normalized sEMG amplitude) during submaximal voluntary isometric contractions at a given relative contraction level (23–25) and its rate of increase with contraction level (23) have been reported to be lower at extended (short muscle) *vs* flexed (long muscle) knee-joint positions. Whilst deficits in these global measures of neuromuscular activation imply attenuated MU recruitment and discharge rate (modulation) at extended knee positions, the effect of knee-joint angle on MU discharge properties remains poorly understood. The only report we are aware of compared vastus lateralis MU discharge rates between different knee-joint angles during the production of consistent absolute torque (25), which, given the profound effect of joint angle on contractile torque production, seems likely to have necessitated differences in neural activation of the muscle and thus precluded any meaningful comparison of neural control.

A potential mechanism for the different modulation of MU discharge rate with knee-joint angle could in part be due to differences in the intrinsic excitability of spinal motoneurons innervating the vastii muscles. The intrinsic excitability of motoneurons is largely determined by the magnitude of persistent inward currents (PICs), which are strong depolarising inward currents facilitated in the dendrites by monoamines (serotonin and noradrenaline) released from the brainstem (26). By amplifying net excitatory synaptic input (up to fivefold), PICs are essential for initiating and sustaining motoneuron discharge, and thus shaping their input-output function (26). Importantly, PICs are highly sensitive to synaptic inhibition, which can attenuate or even terminate their activity (27–29). In decerebrate cat preparations, joint-angle-dependent variations in net inhibitory feedback from muscles surrounding the joint have been shown to modulate the PIC contribution to motoneuron firing (5). Similarly, in humans, greater PIC estimates have been observed during isometric contractions at short compared to long muscle lengths in the tibialis anterior and gastrocnemius medialis (4). Whether similar modulation of PICs occurs in the quadriceps muscles remains unexplored. Furthermore, while PIC magnitude in both ankle and knee muscles has been shown to increase with contraction level (30), it remains unknown whether the magnitude of contraction-level-dependent PIC modulation varies with knee-joint angle.

Therefore, this study aimed to assess the effect of knee-joint angle and contraction level on both the MU discharge rates and the contribution of PICs to MU discharge in the vastii muscles. Using high-density surface electromyography and signal decomposition techniques, MU discharge properties were examined during voluntary isometric knee extension contractions at maximal and several submaximal contraction levels across three knee-joint angles (25°, 55°, and 85° of flexion). We hypothesised that MU discharge rates and their modulation with increasing contraction levels would vary between knee-joint angles, with higher discharge rates and a steeper increase in discharge rate at flexed (i.e., long muscle length) compared to extended (i.e., short muscle length) angles. Furthermore, we hypothesised that modulation of PIC magnitude would mirror the angle-dependent adjustments in MU discharge behaviour.

## MATERIALS AND METHODS

### Participants

Twelve healthy adults (1 female; age, 23 ± 3 years; body mass, 75 ± 11 kg; height, 1.77 ± 0.09 m) participated in this study after providing written informed consent. Participants were asymptomatic and reported no history of musculoskeletal or neuromuscular conditions affecting function of the lower limbs, no history of cardiovascular conditions, and were not consuming any medication known to alter function of the neuromuscular system. The study was approved by Loughborough University Ethical Advisory Committee (Reference ID: 2021-6330-7544) and was conducted in accordance with the Declaration of Helsinki (apart from registration in database).

### Experimental design

Participants visited the laboratory on four occasions separated by 2-7 days. During the first session, participants were familiarised with experimental procedures and practiced experimental tasks performed in subsequent sessions. Then, three experimental sessions were conducted at a consistent time of day (±1 h), with participants instructed to arrive rested and hydrated, and having avoided strenuous exercise and alcohol consumption for 36 h, and caffeine intake for 12 h prior to arrival. All sessions involved unilateral isometric knee-extension contractions, with participants seated on a rigid, custom-built knee-extension dynamometer. The hip-joint angle was fixed at ∼55° of flexion (0° = anatomical position). The dynamometer configuration was adjusted to each participant’s size and kept constant across the sessions. In each experimental session, the neuromuscular assessment was conducted at a different knee-joint angle (∼25, ∼55, or ∼85° of flexion) using an identical experimental protocol. Both legs were tested on each occasion, with a ∼15 min break between the assessments where participants could freely move around the laboratory. The orders of legs (dominant, nondominant) and knee-joint angles (i.e. sessions) tested were randomised. The dominant leg was determined using the lateral preference inventory (31).

### Experimental protocol

Participants first completed a brief warm-up involving a series of submaximal isometric contractions at 50% (× 3), 75% (× 2) and 90% (× 1) of perceived maximal effort (∼4 s). Then, participants performed five MVCs separated by at least 30 s of rest, with twitches superimposed on the final three MVCs to assess voluntary activation. Participants were instructed to ‘push as hard as possible’ for 3-4 s. To facilitate maximal force production, strong verbal encouragement was provided, and a real-time force trace was displayed on a computer monitor ∼1 m in front of the participant with a horizontal cursor indicating the participant’s current highest force. The highest instantaneous voluntary force attained across all MVCs was recorded as the maximal voluntary force (MVF). After ∼3 min of rest, participants performed triangular contractions to 30 and 50% MVF (× 2 at each contraction level) in a randomised order. Each contraction involved a 10-s linear increase and a 10-s linear decrease in force output (rates of force increase/decrease of 3 and 5% MVF/s, respectively). Then, trapezoidal contractions were performed at 10, 30, 50, and 70% MVF (× 2 at each contraction level). The order of the 10-50% MVF contractions was randomised, whereas the 70% MVF contractions were always performed last to minimise decrements in muscle contractile function. Each contraction involved a linear increase in force output to the target level (10% MVF/s), a 10-s plateau, and a linear decrease in force output (10% MVF/s). All submaximal contractions were performed by tracking a triangular/trapezoidal waveform displayed on the computer monitor. Before each contraction, participants were instructed to track the target force trace ‘as accurately as possible’. Between successive trials, sufficient rest was ensured to minimise the potential confounding effects on MU discharge properties (i.e., accommodation of MU recruitment threshold; (32)), with a minimum of 30, 45, and 90 s separating contractions performed at 10-30, 50, and 70% MVF, respectively.

### Experimental procedures

#### Force recordings

Knee-extensor force production was measured using a calibrated S-beam strain gauge (linear range ±1.5 kN; Force Logic, Swallowfield, UK) attached perpendicular and posterior to the participant’s tibia using a bespoke non-extendable webbing strap (35 mm width) tightly fastened ∼3-4 cm superior to the lateral malleolus. Participants were also firmly strapped across the chest and hips to restrict compensatory movement. The analogue force signal was sampled at 2048 Hz and amplified (× 370) using an A/D converter (Micro 1401-4; CED Ltd., Cambridge, UK), and recorded using the Spike2 software (version 10.14; CED Ltd.). The analogue force signal was also concurrently sampled via a 16-bit multichannel amplifier (Quattrocento; OT Bioelettronica, Torino, Italy) and recorded in OTBioLab+ software (version 1.5.7.3; OT Bioelettronica, Torino, Italy) for synchronisation with electromyographic recordings.

#### High-density surface electromyography

Myoelectric activity of vastus lateralis (VL) and vastus medialis (VM) muscles during the first two MVCs (i.e. without a superimposed electrical stimulus) and all submaximal contractions were recorded using two semi-disposable 64-electrode grids (5 columns × 13 rows; interelectrode distance: 8 mm; OT Bioelettronica, Torino, Italy). Following skin preparation involving shaving, light abrasion, and cleansing (Nuprep; Weaver and Company, CO, USA), the grids filled with a conductive paste (Ten20; Weaver and Company) were positioned over the VL and VM muscle bellies in line with the presumed muscle fibre orientation and attached to the skin using disposable bi-adhesive foam layers (SpesMedica, Battipaglia, Italy). Specifically, the VL electrode grid was positioned at an angle of ∼20° relative to the line between the anterior superior iliac spine (ASIS) and the lateral border of the patella, with channel row 6/7 aligned with ∼50% of the thigh length (distance between the knee-joint space and the greater trochanter). The VM electrode grid, however, was placed at an angle of ∼50° relative to the line between the ASIS and the medial border of the patella, with the 4^th^ distal row of electrodes positioned at ∼95% of the line length from the ASIS. Two self-adhesive reference electrodes (3M 2239, Red Dot, Bracknell, UK) were attached over the patella of both legs and two damp strap electrodes (WS2, OT Bioelettronica, Torino, Italy) were fastened in parallel superior to the lateral malleolus of the non-tested leg to ground the signal. High-density sEMG signals were recorded in a monopolar configuration during each contraction. The signals were sampled at 2048 Hz and amplified (× 150) using a multichannel amplifier (16 bit; Quattrocento, OT Bioelettronica, Torino, Italy), band-pass filtered (10-500 Hz), and recorded in OTBioLab+ software (version 1.5.7.3; OT Bioelettronica, Torino, Italy).

#### Percutaneous femoral nerve stimulation

To assess maximal quadriceps voluntary activation and potentiated resting twitch torque, single supramaximal electrical stimuli (square-wave, 200 μs, variable-voltage; DS7AH, Digitimer, Welwyn Garden City, UK) were manually delivered to the femoral nerve during the force plateau and ∼2 s after each of the final three MVCs. Circular self-adhesive gel-coated cathode and anode (32 mm^2^; CF3200, Nidd Valley Medical, Bordon, UK) were placed in the femoral triangle and over the greater trochanter, respectively. The stimulation current was calibrated before the warm-up knee-extension contractions by gradually increasing the stimulus intensity from 20 mA in 40-mA increments until a plateau in the resting quadriceps twitch force was attained. To ensure supramaximal stimulation, 150% of the plateau current was used for assessment (485 ± 246 mA).

#### Knee-joint angle recordings

To measure the knee-joint angle during the isometric knee-extension contractions, sagittal plane video of the participant’s right leg was recorded during the second MVC using a video camera (Panasonic HC-V160; Secaucus, NJ, USA) positioned 3.4 m perpendicular to the participant’s sagittal plane. For the offline analysis, participants had visible markers drawn on the greater trochanter, knee-joint space, and lateral malleolus.

### Data analysis

#### Voluntary and evoked torque

Offline analysis of the force signal recorded during maximal voluntary and evoked contractions was performed in Spike2 (version 10.14; CED, Cambridge, UK). Prior to analysis, voltage force signal was low-pass filtered at 20 Hz (4^th^ order, zero-lag Butterworth filter), converted to force (N), corrected for gravity (subtracting baseline force level), and converted to torque (Nm) via multiplication by lever length (distance between the knee-joint space and the centre of the ankle strap). The maximal voluntary torque (MVT) was measured as the peak instantaneous voluntary torque value recorded across the five MVCs. For each of the three MVCs involving percutaneous electrical stimulation, the amplitudes of the superimposed (SIT) and potentiated resting (Q_tw_) twitches were measured as a difference in torque from twitch onset to peak, with the values averaged and the average Q_tw_ also normalised to MVT. Quadriceps voluntary activation level during each of the final three MVCs was estimated as the ratio of the SIT and Q_tw_ amplitudes subtracted from complete muscle activation using the classic equation, 1-SIT/Q_tw_ × 100 (18). If torque at stimulus delivery (T_SIT_) was lower than the peak instantaneous voluntary torque attained during the assessed contraction (T_peak_) by > 2.5%, a correction equation was used to account for this deviation: 100 – SIT × [T_SIT_/T_peak_]/Q_tw_ × 100 (33). The maximal recorded quadriceps voluntary activation value was used for further analysis.

#### Motor unit discharge properties

Offline analysis of the HDsEMG data was performed in MATLAB (R2021b; Mathworks Inc., MA, USA).

##### High-density sEMG signal processing

Semiautomatic tool DEMUSE (University of Maribor, Slovenia) was used for signal processing. After the signals were band-pass (20-500 Hz, zero-lag, 5^th^ order Butterworth filter) and notch filtered (50 Hz and its higher harmonics), the channels demonstrating low signal-to-noise ratio and/or presence of artefacts were removed from further processing (≤ 5% of the channels). Then, signals were decomposed into MU discharge timings using the validated Convolution Kernel Compensation algorithm (34).

##### Submaximal voluntary contractions

Signals from each triangular/trapezoidal contraction were decomposed independently (50 decomposition runs). To track the activity of individual MUs across the two triangular/trapezoidal contractions performed at the same contraction level and knee-joint angle, decomposition results were first concatenated, and then MU filters extracted from decomposition of the signal of one contraction were applied to the signal of the other contraction. Duplicates of MUs produced through this procedure (>30% of shared discharge timings; 0.5 ms match tolerance) were removed, with the spike trains of the MU exhibiting lower decomposition accuracy (i.e., lower pulse-to-noise ratio; (35)) discarded. To optimise MU filter estimation accuracy, the identified MU spike trains were visually inspected and manually edited using the previously described iterative procedure (36). Note that MUs were tracked only across the two contractions performed at the same contraction level and the same knee-joint angle but not across different contraction levels or same-level contractions performed at different knee-joint angles.

##### Maximal voluntary contractions

Signals recorded during the two MVCs were concatenated (37, 38) prior to decomposition to enhance decomposition accuracy and facilitate MU identification. As a result, individual MUs were automatically identified in either one or both MVCs (i.e., tracked MUs). Additionally, a second round of decomposition was conducted which, compared to the first round, included the entire recorded frequency band (10-500 Hz) without any notch filtering (50 Hz and its harmonics). Any duplicate MUs identified across the two decomposition rounds (30 decomposition runs each) were removed, and the remaining MU spike trains were examined and manually edited (see *Submaximal voluntary contractions* for details).

Data extraction was performed using custom-written MATLAB scripts. Only MUs exhibiting pulse-to-noise ratio ≥ 28 dB and regular discharge patterns (all interspike intervals < 0.5 s and a minimum of 40, 10, and 20 discharges for trapezoidal, triangular, and maximal contractions, respectively) were kept for further analysis.

##### Motor unit discharge rate at different submaximal contraction levels and MVCs

Steady-state discharge rate of individual MUs identified during trapezoidal contractions or MVCs was quantified from the decomposed binary spike trains. The instantaneous MU discharge rate (DR) was first computed as an inverse of the interspike intervals. Interspike intervals < 15 ms or > 450 ms were excluded from the analysis. For individual MUs identified during *submaximal trapezoidal contractions (10-70% MVT)*, the mean discharge rate was calculated as the average across the 10-s plateau at the target torque. Conversely, for individual MUs identified during *MVCs*, a rolling average of instantaneous discharge rates during the torque plateau was first computed for every window of 10 discharges with a 1 discharge shift. The maximal value was noted as the discharge rate during MVC (DR_MVC_). Only trials during which peak instantaneous torque attained was ≥ 85% MVT were included in further analysis.

The recruitment threshold of individual MU identified during submaximal trapezoidal contractions was calculated as the mean torque corresponding to the first two discharge timings and was normalised to MVT. In contrast, for MU identified during MVCs, recruitment threshold could not be reliably determined due to relatively rapid and variable (within and between participants) rate of torque development.

#### Intrinsic electrical properties of motoneurons

##### Smoothing of the instantaneous motor unit discharge rates

Analyses of intrinsic electrical properties of motoneurons were conducted using MU spike trains identified during triangular contractions (30 and 50% MVT). Prior to analyses, the decomposed binary MU spike trains were smoothed using support vector regression (SVR) as described previously (39). In short, continuous estimates of discharge rates from MU recruitment to derecruitment were generated using an SVR model trained on the instantaneous MU discharge rates and fitted using parameters suggested previously (40).

##### Paired motor unit analysis

To estimate the magnitude of PICs we used the established paired MU analysis (41–43), which quantifies the onset-offset discharge rate hysteresis of a higher-threshold (test) MU with respect to a lower-threshold (reporter) MU (41). The reporter MU provides an estimate of the net common synaptic input to the motoneuron pool at recruitment and derecruitment of a higher-threshold (test) MU (41). The difference between the control MU discharge rates at recruitment and derecruitment of the test MU is defined as ΔF (41).

Pairs of test and reporter MUs active during individual contractions were considered suitable if the following criteria were met: (1) the test MU was recruited ≥ 1 s after the reporter MU to ensure that PICs in the reporter MU were fully active before test MU recruitment (44); (2) the paired MUs demonstrated a rate-rate correlation of r > 0.70 to ensure shared synaptic input between the test and reporter MUs (45); and (3) discharge rate of the reporter MU increased by > 0.5 pps while the test MU was active to minimise the impact of reporter MU saturation on ΔF calculation (46). The ΔF of each test MU was calculated as the average of ΔF values across all suitable reporter MU pairs (Figure 1A) (30, 47, 48).

**Figure 1.**
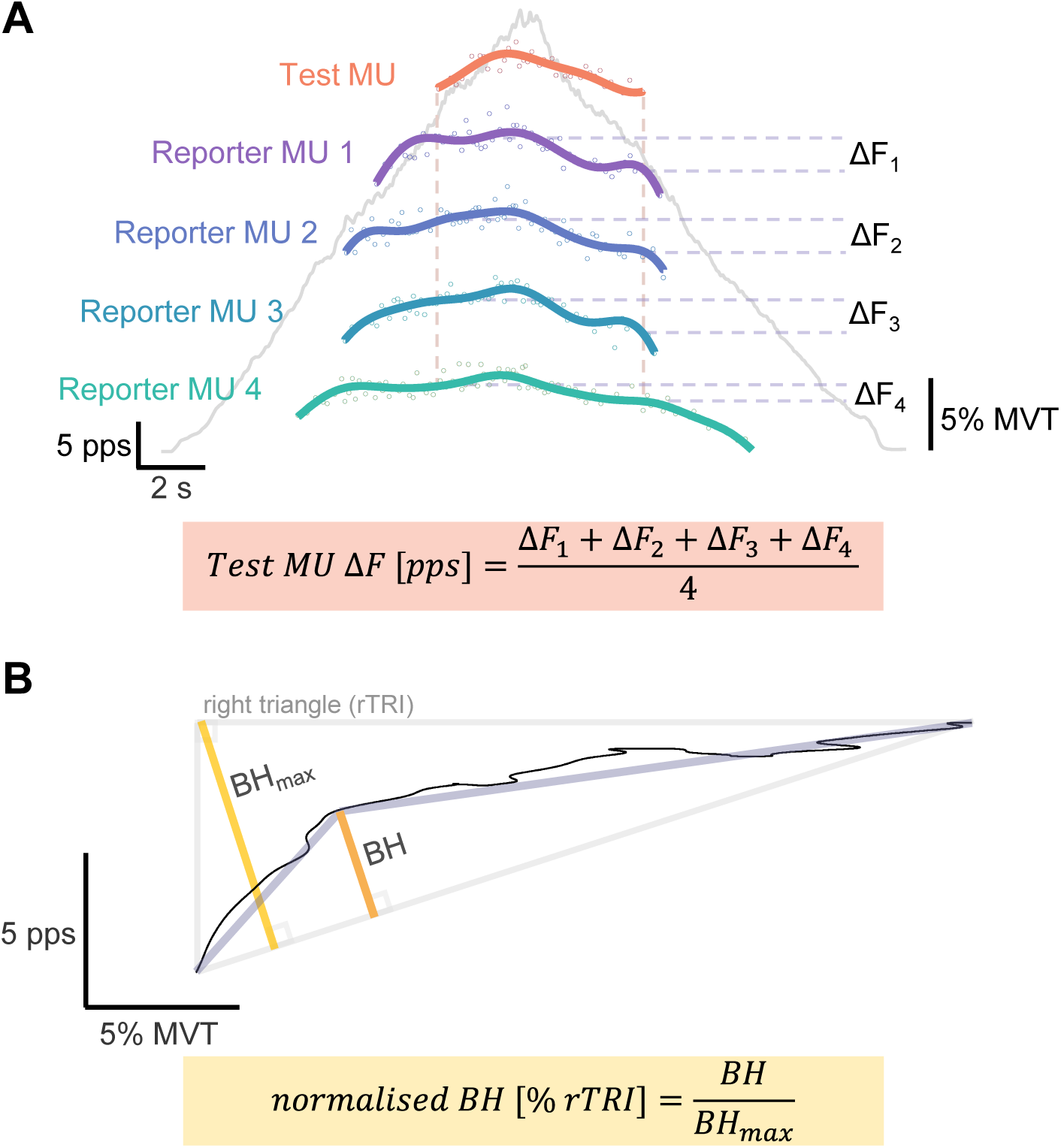
Schematic of ΔF and brace height calculation. Onset-offset discharge rate hysteresis (ΔF; A) of a test motor unit (MU) was calculated as the average of ΔF values calculated for all suitable reporter MU pairs. Degree of MU discharge rate nonlinearity (B) was quantified geometrically as a maximal deviation of discharge rate trajectory from a theoretical linear increase (brace height; BH) and normalised to the BH of the right triangle (rTRI) representing the maximal possible BH (BH_max_). MVT, maximal voluntary torque; pps, pulses per second.

##### Geometric analysis

To quantify the degree of nonlinearity in MU discharge rate with respect to torque output during the ascending phase of the triangular contractions we used the pseudo-geometric analysis (40). In this study, torque output was used as an estimate for the net excitatory synaptic input. For each identified MU, the SVR-smoothed discharge rates were first plotted as a function of torque output during a contraction. Then, a straight line was fitted between the first point (onset discharge rate) and the maximal value of the smoothed discharge rate estimates (i.e., peak discharge rate). This line represented a trace of a theoretical linear increase in discharge rate with torque production. Finally, the magnitude of the maximum orthogonal vector from this straight line to the smoothed discharge rate trace was quantified and represented the brace height. To account for the scaling of the brace height with the range of MU discharge rate modulation during a contraction, the brace height values were normalised to the height of a right triangle with the straight line from the onset and peak discharge rate delimiting the hypothenuse. The right triangle represents the theoretical MU discharge rate profile with full PIC activation and thus maximum possible brace height value (40) (Figure 1B). MUs with the peak discharge rate occurring after the peak torque or with normalised brace height greater than 200% were removed from further analysis. Finally, the recruitment threshold of individual MU was determined as the torque at the timing of the first point (discharge estimate) of the smoothed DR.

#### Knee-joint angle

To measure the knee-joint angle during an MVC, video recordings of the participant’s right leg performing an MVC were digitised and the angle between visible markers on the greater trochanter, knee-joint space, and lateral malleolus was measured using an open-access video-analysis software (Kinovea, version 0.9.5; https://www.kinovea.org/; Figure 2).

**Figure 2.**
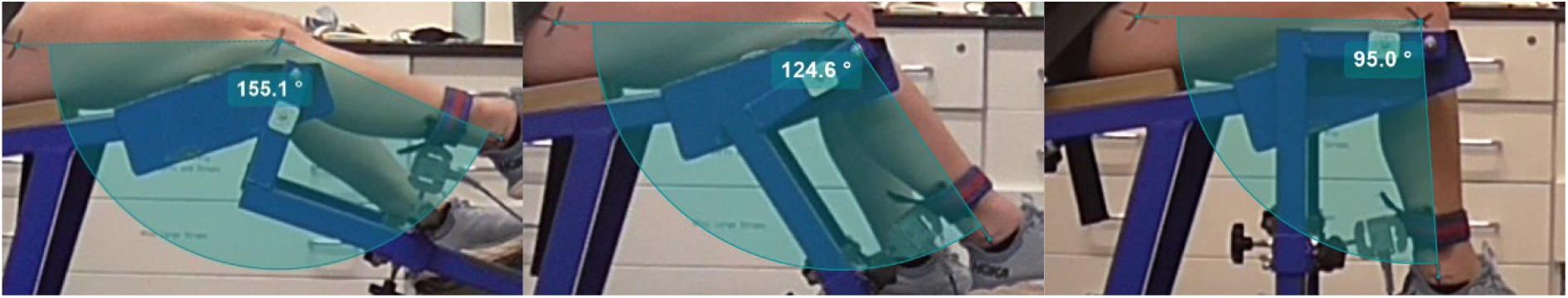
Measurements of the knee-joint angle during maximal voluntary isometric knee-extension contractions. Example sagittal plane images of one participant’s knee during maximal voluntary isometric knee-extension contractions at ∼25° (left), ∼55° (middle), and ∼85° (right) of flexion. The knee-joint angle was measured digitally as the angle between the greater trochanter, knee-joint centre, and lateral malleolus, which was then subtracted from 180° (i.e., full extension) to determine the angle of flexion. The group means and standard deviations of the measured knee-joint angles at the three tested positions were 25 ± 1°, 54 ± 2°, and 83 ± 2° of flexion.

### Statistical analysis

Statistical analysis was performed in RStudio (version 2023.12.1+402). To account for the variability in the outcome variables within and between participants, linear mixed-effect models were constructed (*lme4* package; version 1.1-35.1, (49)). MVT, maximal quadriceps voluntary activation, absolute potentiated resting twitch torque, and normalised potentiated resting twitch torque [% MVT] were compared between the knee-joint angles (25, 55, and 85° of flexion; fixed factor), with individual participants and the leg (dominant, non-dominant) included as random factors. The following model structure was used for the force related outcome variables: outcome variable ∼ 1 + knee-joint angle + (1|participant identifier) + (1|leg). Mean discharge rate during trapezoidal contractions, ΔF, brace height, and onset and peak discharge rate during triangular contractions were compared between the knee-joint angles, contraction levels (mean discharge rate: 10, 30, 50, and 70% MVT; ΔF, brace height, and onset and peak discharge rate: 30 and 50% MVT) and muscles (VL and VM; fixed factors), with MUs nested within individual participants, and with the leg and individual trials included as random factors. As MUs were not tracked across contraction levels or knee-joint angles and thus different samples of MUs were observed in each testing ‘condition’, MUs were given unique names and were nested within the ‘parent’ participant. Additionally, MU recruitment threshold was included in the model as a covariate to control for the effect of the recruitment threshold on MU discharge properties. The following model structure was used: outcome variable ∼ 1+ knee-joint angle × contraction level × muscle + MU recruitment threshold + (1|participant identifier/MU identifier) + (1|leg) + (1|trial identifier). DR_MVC_ was compared between the knee-joint angles and muscles (fixed factors), with MUs nested within individual participants, and the leg and individual trials included in the model as random factors. MU recruitment threshold was not determined for MUs observed during MVCs due to relatively rapid and variable rate of torque development and was thus not included in the model as a covariate. The following model structure was used: discharge rate ∼ 1+ knee-joint angle × muscle + (1|participant identifier/MU identifier) + (1|leg) + (1|trial identifier). After fitting the model, normal distribution and linearity of the residuals were assessed using histograms and quantile-quantile plots, respectively. In cases of non-normal distribution of residuals, data was transformed using either a square root (DR_MVC_, brace height) or log1p (maximal quadriceps voluntary activation, mean DR during trapezoidal contractions) transformation. The transformation function was selected based on a visual inspection of the distribution and linearity of residuals and the goodness of the model fit (Akaike Information Criterion and Bayesian Information Criterion). Outliers were assessed using box plots. There were no cases of outliers negatively affecting linearity and skewing the distribution of residuals, therefore no cases were removed. Type III ANOVA with Satterthwaite’s method was used to test the effect of the fixed factors on the outcome variables. The origin of significant main effects and two-way interactions were further explored using simple and interaction contrasts of estimated marginal means with post-hoc tests (*emmeans* package; version 1.8.9). Bonferroni adjustment was used for multiple comparisons. Significance was set at p < 0.05. Significant three-way interactions (knee-joint angle × contraction level × muscle) were not explored further due to high level of complexity.

## RESULTS

### Knee-joint angle

On average, the knee-joint angles measured with sagittal plane video during maximal voluntary contractions at the three tested positions were 25 ± 1°, 54 ± 2°, and 83 ± 2° of flexion.

### Number of identified motor units

Decomposition of high-density sEMG signals recorded from VL and VM muscles during trapezoidal, triangular and maximal voluntary isometric knee-extension contractions yielded a total of 8962, 3529, and 1127 MUs, respectively. Of those 94.3%, 90.6%, and 79.3% MUs identified during trapezoidal, triangular, and maximal contractions, respectively, were tracked across both contractions performed at the same contraction level within each recording condition (as defined by the leg, knee-joint angle and muscle). Table 1 presents total numbers of MUs identified across all recorded contractions for each leg (dominant, nondominant) and each knee-joint angle, as well as participant averages and standard deviations per contraction. As two contractions were performed in each recording condition (defined by the leg, knee-joint angle, muscle, and contraction level), with MUs tracked across the two contractions, Table 1 also presents numbers of unique MUs identified per recording condition.

**Table 1.** The number of identified motor units with high-density surface EMG decomposition per leg and per knee-joint angle.

| Number of identified motor units |  |  |  |  |  |  |  |  |
| --- | --- | --- | --- | --- | --- | --- | --- | --- |
|  |  |  | Trapezoidal contractions |  | Triangular contractions |  | Maximal contractions |  |
|  |  |  | All | Per condition | All | Per condition | All | Per condition |
| Leg | Nondominant | Total | 4518 | 2391 | 1737 | 962 | 463 | 301 |
| | | Mean ± SD | 8.1 ± 4.1 | 8.6 ± 4.2 | 6.4 ± 3.7 | 7.0 ± 4.0 | 4.6 ± 3.2 | 4.6 ± 3.2 |
|  | Dominant | Total | 4444 | 2345 | 1792 | 968 | 664 | 379 |
| | | Mean ± SD | 7.9 ± 4.2 | 8.3 ± 4.3 | 6.5 ± 3.1 | 7.0 ± 3.4 | 5.9 ± 4.5 | 5.9 ± 4.4 |
| Knee-joint angle | 25° | Total | 3314 | 1770 | 1367 | 744 | 464 | 285 |
| | | Mean ± SD | 8.9 ± 3.9 | 9.5 ± 4.0 | 7.4 ± 3.8 | 8.0 ± 4.1 | 6.9 ± 4.8 | 7.1 ± 5.2 |
|  | 55° | Total | 2990 | 1573 | 1111 | 615 | 385 | 225 |
| | | Mean ± SD | 7.9 ± 4.3 | 8.3 ± 4.5 | 6.0 ± 3.3 | 6.5 ± 3.7 | 5.4 ± 3.4 | 5.5 ± 3.4 |
|  | 85° | Total | 2658 | 1393 | 1051 | 571 | 278 | 170 |
| | | Mean ± SD | 7.1 ± 4.5 | 7.4 ± 4.7 | 5.8 ± 3.6 | 6.3 ± 3.9 | 4.0 ± 3.8 | 4.0 ± 3.5 |

Note: 'All' represents the number of all motor units (MUs) identified across all recorded contractions; 'Per condition' represents the number of unique MUs identified across two contractions performed at the same contraction level per recording condition (as defined by the leg, knee-joint angle and muscle). 'Total' represents the number of all identified MUs across all contractions of each type across all participants. Means and SD were calculated per participant per contraction; values were averaged across knee-joint angle, muscle, and contraction level. SD, standard deviation.

### Global measures of neuromuscular function

Main effects of the knee-joint angle were found for isometric MVT (F_2,57_ = 165.0; p < 0.0001), quadriceps voluntary activation (F_2,56_ = 17.1; p < 0.0001), as well as absolute (F_2,57_ = 167.5; p < 0.0001) and normalised quadriceps potentiated twitch torque (F_2,57_ = 39.3; p < 0.0001). MVT differed between all the knee-joint angles tested (all p < 0.0001), whereby it was the greatest at the middle knee-joint angle (55°) and the smallest at the extended knee-joint angle (25°; Figure 3A). Maximal voluntary quadriceps activation was the lowest at the middle knee-joint angle (55°; *vs* 25°: p < 0.0001, *vs* 85°: p = 0.0003), with no difference between the two extreme positions (25° and 85°; p = 0.4780; Figure 3B). Absolute quadriceps potentiated twitch torque differed between all the knee-joint angles tested (p ≤ 0.0006; Figure 3C), whereby it was the greatest at the middle (55°), and the smallest at the extended (25°) position. When expressed as a proportion of MVT, quadriceps potentiated twitch torque was the greatest at the most flexed knee-joint angle (85°; *vs* 25° and 55°: p < 0.0001), with no difference between the two more extended knee-joint angles (25° and 55°; p = 0.0944; Figure 3D).

**Figure 3.**
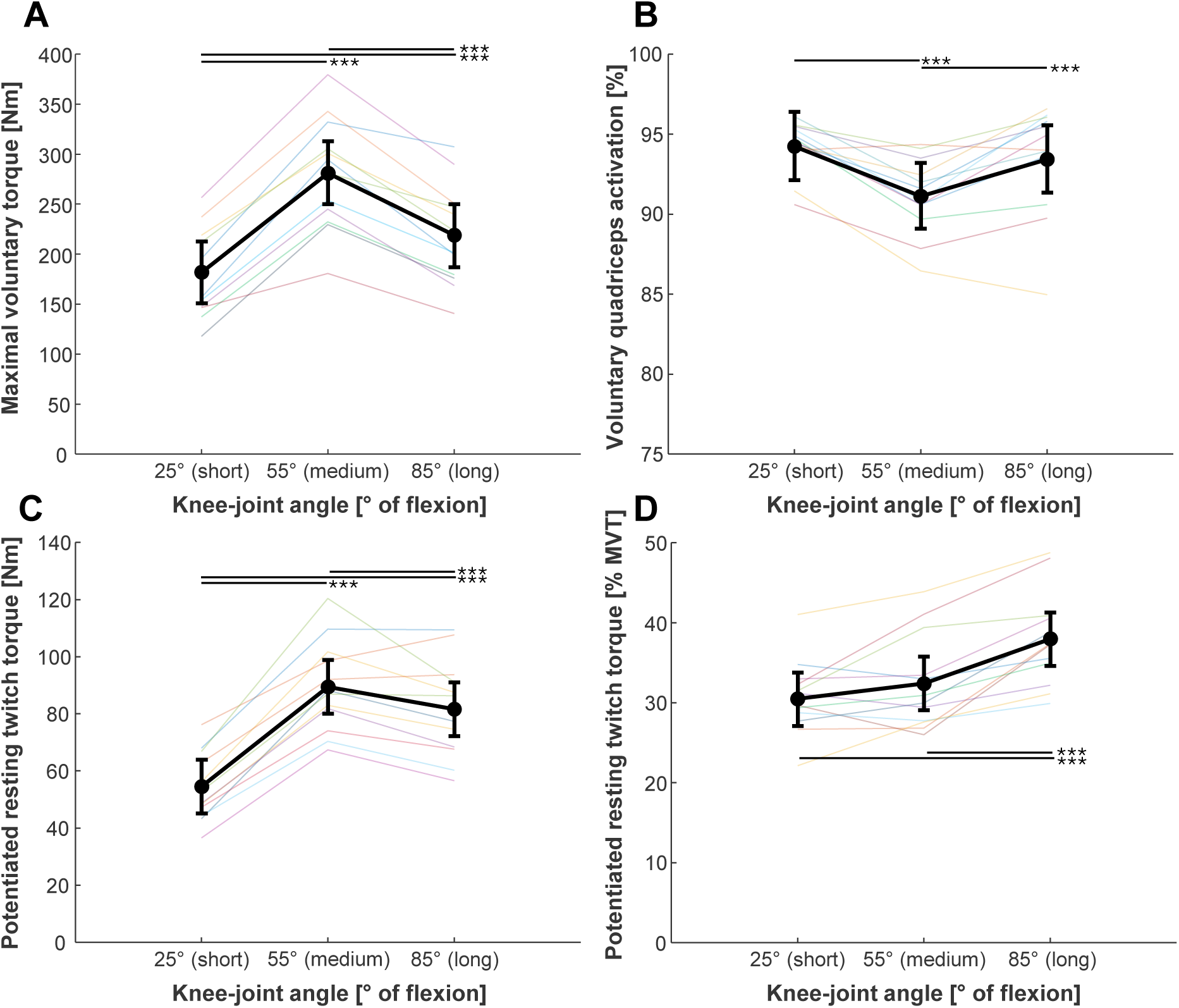
Measures of global neuromuscular function of the knee extensors at different knee-joint angles. Maximal voluntary torque (MVT; A), voluntary quadriceps activation level during maximal voluntary contractions (B), absolute potentiated quadriceps resting twitch torque (C), potentiated quadriceps resting twitch torque normalised to MVT (D) at 25° (short), 55° (medium), and 85° (long muscle length). Black, filled circles represent the estimated marginal means of the group, with error bars representing 95% confidence intervals. Thin lines of different colours represent individual participants’ data. ***, p < 0.001.

### Motor unit discharge rate modulation with contraction level

#### Submaximal voluntary isometric knee-extension contractions

Mean MU DR during the plateau phase of trapezoidal contractions showed main effects of the knee-joint angle (F_2,4560_ = 106.2; p < 0.0001), contraction level (F_3,5419_ = 3296.4; p < 0.0001), and muscle (F_1,4520_ = 49.2; p < 0.0001), as well as interaction effects between angle and contraction level (F_6,4529_ = 11.7; p < 0.0001; Figure 4A), and angle and muscle (F_2,4519_ = 4.6; p = 0.0105). The overall mean MU DR was lower at 55° of flexion (middle) than at 25 and 85° (both p < 0.0001), with no difference between the two extreme angles (p = 1.0000). These pairwise differences, however, depended on the contraction level. Specifically, when compared to the two extreme angles (i.e., 25 and 85°), MU DR at 55° was lower at all contraction levels >10% MVT (all p < 0.0001). At 10% MVT, however, DR at 55° was only lower than 25° (p = 0.0015), but not 85° (p = 1.000). Compared to 25°, DR at 85° was lower at 10 and 30% (p ≤ 0.0375), similar at 50% (p = 1.0000) and higher at 70% MVT (p < 0.0001). As expected, MU DR increased with contraction level, with similar increases at all knee-joint angles up to 50% MVT. However, when contraction level increased from 50 to 70% MVT, there was a greater upregulation in MU DR at the most flexed angle (85°) compared to 55° or 25° (p ≤ 0.0009), but with no difference between 55 and 25° (p = 1.0000). When comparing the vastii muscles, the overall mean MU DR of the VM was higher than that of the VL (11.4 [10.8, 11.9] pps *vs* 11.1 [10.5, 11.6]; p < 0.0001). Considering individual angles, VM MU DR was higher than the VL at 25° (11.7 [11.2, 12.3] pps *vs* 11.3 [10.7, 11.8]; p < 0.0001) and 85° (11.8 [11.2, 12.3] pps *vs* 11.2 [10.7, 11.8]; p < 0.0001), but similar at 55° (10.8 [10.2, 11.3] pps *vs* 10.7 [10.1, 11.2]; p = 0.0844).

**Figure 4.**
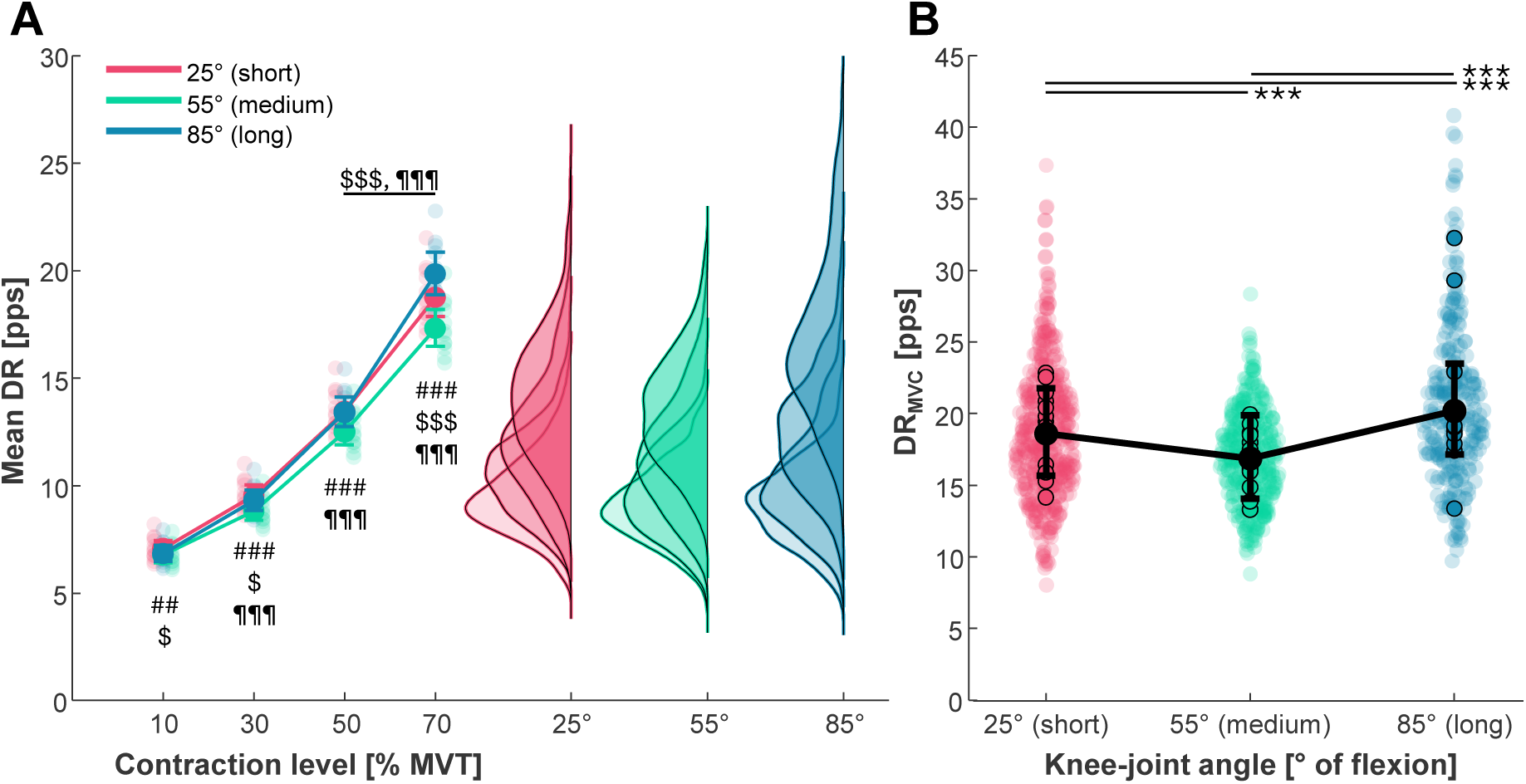
Motor unit discharge rate as a function of contraction level and joint angle. Motor unit discharge rate of the vastii muscles (vastus lateralis and medialis) during trapezoidal isometric knee-extension contractions at 10, 30, 50, and 70% of maximal voluntary torque (MVT; A) and during maximal voluntary contractions (DR_MVC_; B) at 25° (short), 55° (medium), and 85° (long muscle length) of knee flexion. For illustrative purposes, the data are grouped for both vastii muscles. A: The larger, opaque circles linked with a solid line represent estimated marginal means of the group as calculated from the linear mixed-effect statistical model, with the error bars representing the 95% confidence intervals. The smaller, transparent circles represent individual participant estimated marginals. The kernel density plots depict distribution of individual motor unit data points. The horizontal black line denotes statistical results of interaction contrasts (knee-joint angle × contraction level). #, 25 *vs* 55°; $, 25 *vs* 85°; ¶, 55 *vs* 85°. E.g., #, p < 0.05; ##, p < 0.01; ###, p < 0.001. B: Black, filled circles represent the estimated marginal means of the group as calculated from the linear mixed-effect statistical model, with error bars representing 95% confidence intervals. The smaller, opaque circles represent individual participants estimated marginals and the small, transparent circles represent individual motor unit data points. ***, p < 0.001.

#### Maximal voluntary isometric knee-extension contractions

There were main effects of the knee-joint angle (F_2,663_ = 40.1; p < 0.0001; Figure 4B) and muscle (F_1,666_ = 4.9; p = 0.0278) on MU DR during MVCs. DR_MVC_ differed between all the knee-joint angles (p ≤ 0.0001), being the lowest at 55° (16.8 [14.1, 19.9] pps) and the highest at 85° (20.2 [17.1, 23.5] pps), with 25° (18.6 [15.7, 21.8] pps) in-between. When comparing the two vastii muscles, VM DR_MVC_ was higher than VL (18.8 [16.2, 21.7] *vs* 18.1 [15.5, 21.1] pps; p = 0.0280).

### Intrinsic electrical properties of motoneurons

#### Onset-offset motor unit discharge rate hysteresis (ΔF)

ΔF showed main effects of knee-joint angle (F_2,1377_ = 111.3; p < 0.0001), contraction level (F_1,1527_ = 47.0; p < 0.0001), and an angle × contraction level interaction (F_2,1374_ = 7.3; p = 0.0007; Figure 5A). Overall, ΔF decreased with the degree of knee flexion (p ≤ 0.0005). Similarly, when considering individual contraction levels, at 30% MVT, ΔF differed between all angles in the order of 25° > 55° > 85° (all p < 0.0001), whereas, at 50% MVT, ΔF was higher at 25° than the other two angles (both p < 0.0001), which were similar (p = 1.0000). ΔF also increased with contraction level (p < 0.0001), with the magnitude of the contraction-level-dependent increase (all p ≤ 0.0026) being greater at 85° than at 25° and 55° (interaction contrasts, p ≤ 0.0428), with no difference between 25° and 55° (interaction contrast, p = 0.8108).

**Figure 5.**
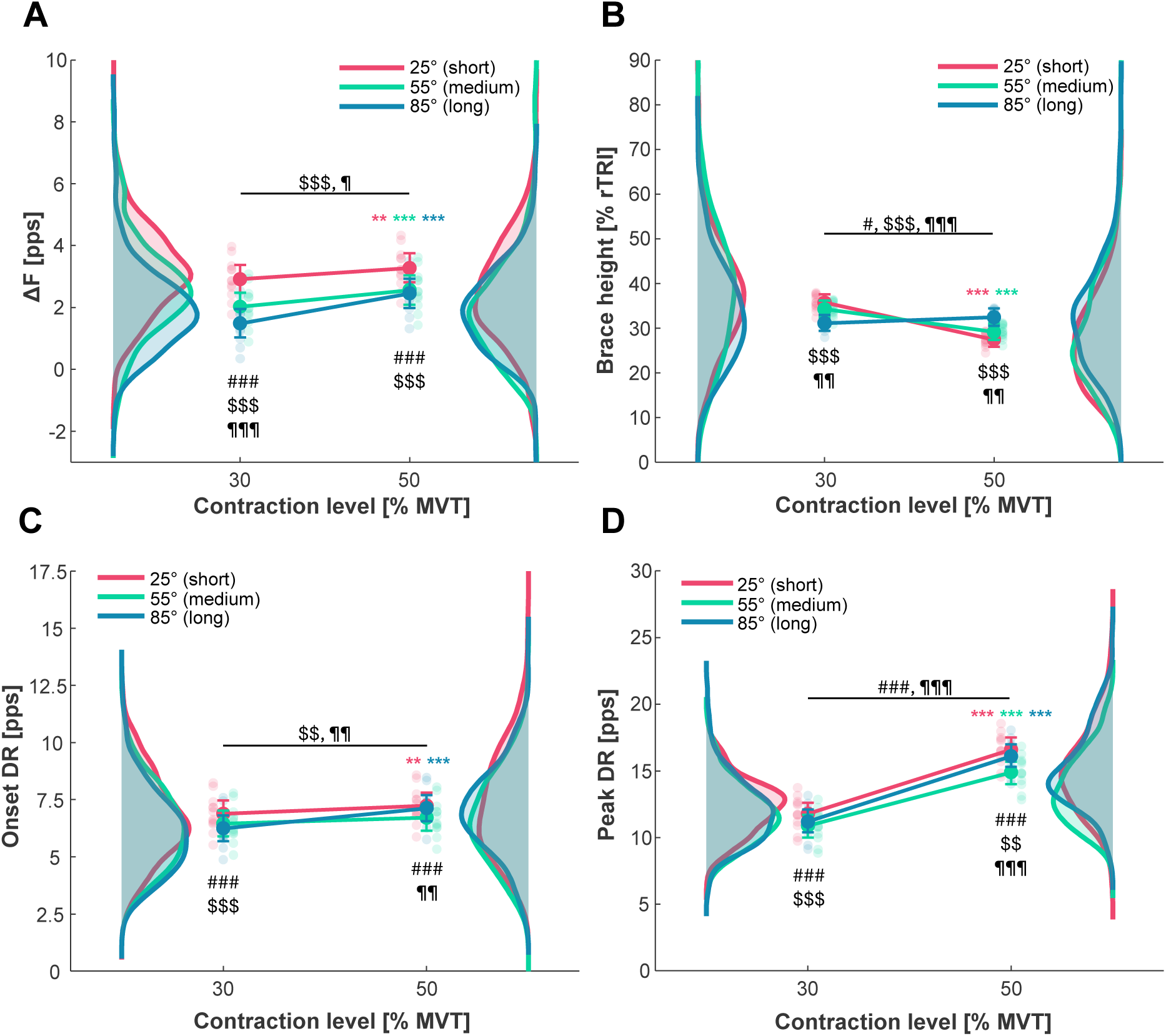
Discharge properties of the vastii motoneurons relative to the contraction level and knee-joint angle. ΔF (A) as a measure of onset-offset discharge rate (DR) hysteresis, brace height (B) as a measure of input-output nonlinearity (ascending phase), onset discharge rate (C) and peak discharge rate (D) of the vastii muscles (vastus lateralis and medialis) motor units during triangular isometric knee-extension contractions up to 30 and 50% of maximal voluntary torque (MVT) and at 25° (short), 55° (medium), and 85° (long muscle length) of knee-joint flexion. For illustrative purposes, the data are grouped for both vastii muscles. The larger, opaque circles linked with a solid line represent estimated marginal means of the group as calculated from the linear mixed-effect statistical model, with the error bars representing the 95% confidence intervals. The smaller, transparent circles represent individual participant estimated marginals. The kernel density plots depict distribution of individual motor unit data points. 50 *vs* 30% MVT: *, p < 0.05; **, p < 0.01; ***, p < 0.001. #, 25 *vs* 55° of knee flexion; $, 25 *vs* 85° of knee flexion; ¶, 55 *vs* 85° of knee flexion. E.g., #, p < 0.05; ##, p < 0.01; ###, p < 0.001. The horizontal black line denotes statistical results of interaction contrasts (knee-joint angle × contraction level).

#### Degree of motoneuron input-output nonlinearity (brace height)

For brace height, there was no main effect of the knee-joint angle (F_2,1603_ = 0.1; p = 0.8740), but a main effect of contraction level (F_1,1640_ = 34.6; p < 0.0001), as well as knee-joint angle × contraction level (F_2,1600_ = 27.8; p < 0.0001; Figure 5B) and muscle × contraction level (F_1,1594_ = 8.1; p = 0.0045) interactions. Simple contrasts revealed that at both 25° and 55° brace height decreased with an increase in contraction level (both p < 0.0001), whereas at 85° it remained unchanged (p = 0.2137). The difference in the responses was reinforced by interaction contrasts, which revealed a progressively smaller contraction-level-dependent change between all three angles in the order of 25° > 55° > 85° (p ≤ 0.0325). When considering individual contraction levels, brace height at 30% MVT, was lower at 85° (p ≤ 0.0024), whereas at 50% MVT it was higher at 85° (p ≤ 0.0045) compared to 25° and 55°, with no difference between 25° and 55° at either contraction level (p ≥ 0.1755). Finally, with contraction level, brace height generally decreased more in VL than in VM (−5.6 *vs* −2.5% rTRI, respectively).

#### Onset motor unit discharge rate

Onset DR revealed main effects of the knee-joint angle (F_2,1881_ = 18.9; p < 0.0001) and contraction level (F_1,1937_ = 32.2; p < 0.0001), and a knee-joint angle × contraction level interaction (F_2,1877_ = 7.5; p = 0.0006; Figure 5C). Whilst the overall onset DR was higher at 25° than at 55° and 85° (both p < 0.0007), with no difference between 55° and 85° (p = 0.7000), these between-angle differences depended on contraction level. At 30% MVT, onset DR was higher at 25° than at 55° and 85° (p ≤ 0.0002), with no difference between the two more flexed positions (p = 0.2235). In contrast, at 50% MVT, onset DR was higher for 25° and 85° than 55° (p ≤ 0.0059), with no difference between the two extreme positions (p = 1.0000). On average, onset DR increased with contraction level, but in a joint-angle-dependent manner. Specifically, simple contrasts showed that at 25° and 85° onset DR increased (p ≤ 0.0054), whereas at 55° it remained unchanged (p = 0.0587). Interaction contrasts further revealed that the contraction-level-dependent increase in onset DR was greater at 85° than at 25 and 55° (p ≤ 0.0037), with no difference in the magnitude of the responses between 25° and 55° (p = 1.0000).

#### Peak motor unit discharge rate

For peak DR, there were main effects of knee-joint angle (F_2,1911_ = 85.8; p < 0.0001), contraction level (F_1,2188_ = 2021.1; p < 0.0001), and muscle (F_1,1917_ = 23.7; p < 0.0001), as well as a knee-joint angle × contraction level interaction (F_2,1908_ = 12.8; p < 0.0001; Figure 5D). On average, peak DR differed between all angles in the order of 55° < 85° < 25° (12.9 [11.8, 13.9] *vs* 13.7 [12.6, 14.8] *vs* 14.2 [13.1, 15.3] pps, respectively; all p < 0.0001). Within individual contraction levels, peak DR at 30% MVT was higher at 25° than at 55° and 85° (p ≤ 0.0001), with no difference between 55° and 85° (p = 0.0525). In contrast, at 50% MVT, peak DR was significantly different between all three angles in the order 55° < 85° < 25° (p ≤ 0.0099). With contraction level, peak DR increased (all p < 0.0001) in a joint-angle-dependent manner, with a lower gain at 55° than 25° and 85° (both p < 0.0001), but no difference between 25° and 85° (p = 1.0000). On average, peak DR was higher for the VM than VL (13.4 [12.4, 14.4] *vs* 13.8 [12.8, 14.8] pps, respectively; p < 0.0001).

## DISCUSSION

This study examined the effect of the knee-joint angle (25° *vs* 55° *vs* 85° of flexion) on MU discharge properties of the vastii muscles and their modulation with contraction level (% MVT at the given knee position). As hypothesised, motoneuron output to the vastii and its increase with contraction level was different between the three tested knee-joint angles, which was accompanied by differential modulation of motoneuron intrinsic electrical properties across the positions. Specifically, both maximal voluntary quadriceps activation and vastii MU discharge rates (maximal and submaximal) were the lowest at the mid-range knee-joint position (55°; medium muscle length), at which the contractile and maximal voluntary torque production capacities were the greatest. Moreover, the greatest MU discharge rate during high force levels (≥70% MVT) was observed at the most flexed position (85°; long muscle length), stemming from a greater discharge rate upregulation required to increase torque output between 50 and 70% MVT compared to the two more extended knee positions (medium and short muscle lengths). While the onset-offset MU discharge rate hysteresis (ΔF) decreased and the degree of input-output nonlinearity (brace height) did not vary as a function of knee flexion (increasing muscle length), the most flexed position was also distinct for a greater increase in the MU discharge rate hysteresis and a lesser decline in the input-output nonlinearity as a function of contraction level when compared to the two more extended positions. This suggests a greater gain in motoneuron excitability with increasing net excitatory drive when the knee is flexed, and the muscle is at a longer length.

### Vastii motor unit discharge rates as a function of knee joint angle and contraction level

In this study, a ‘U’ shaped relationship was observed between voluntary quadriceps activation and the knee-joint angle, with the greatest deficit in activation noted in the middle knee-joint position and no differences found between the more extended and flexed knee positions. These findings are consistent with those by Suter & Herzog (50), and suggest the greatest level of quadriceps inhibition occurs at the mid-range knee position. Notably, however, whilst previous studies have generally reported lower voluntary activation at the mid-range compared to flexed knee positions (19, 20, 51), the findings at extended positions have been mixed; some studies found maximal quadriceps activation level to be lower (19), similar (20, 21), or higher (52, 53) to that at mid-range positions. Although the nature of these discrepant findings is unclear, in this study, the results on the whole-muscle level were reinforced by adjustments in the activity of individual MUs, whereby the pattern of MU discharge rate during maximal voluntary isometric contractions as a function of knee-joint angle followed the general pattern of quadriceps activation. Therefore, we demonstrate, for the first time, that knee-joint-angle-dependent variations in maximal voluntary activation are underpinned by corresponding adjustments in motoneuronal output to the vastii muscles.

In the current study, we found MU discharge rate at contraction levels >10% MVT to be lowest at the mid-range position (on average ∼7% lower compared to extended and ∼8% lower compared to flexed position). Whereas at high contraction levels (≥70% MVT) the discharge rate was higher at the most flexed knee position (long muscle length) than the two more extended positions, with the largest differences compared to the mid-range position (higher by ∼15% at 70% MVT and by ∼20% during MVC). Despite the functional importance of the quadriceps to human mobility and locomotion, only one previous study has examined the effect of knee-joint angle on MU discharge rate of the vastii muscles (25). These authors, however, chose to compare MU discharge rates at the same *absolute* submaximal torque levels and studied a relatively small number of MUs (N = 46) obtained with intramuscular EMG, which makes their results difficult to compare with the present study. In other muscles, however, mixed findings have been reported, including similar MU discharge rate at all muscle lengths/joint angles (*tibialis anterior:* (6, 7); *triceps brachii:* (8)), higher at short (*biceps brachii:* (9); *hamstrings:* (10)) or higher at long (*triceps surae:* (11–13)). These inconsistent findings may be attributed to methodological factors (e.g., small samples of MUs sampled with intramuscular EMG, differing contraction levels, varied statistical approaches). Alternatively, it is possible that joint-angle-induced adjustments in neural control of muscle torque production is, at least in part, muscle- or joint-dependent. For example, unlike in the knee extensors, voluntary activation level has been shown not to vary as a function of joint angle in the elbow flexors and extensors (54),or ankle dorsiflexors (6).

There are several potential explanations for the greater degree of vastii muscle inhibition and/or disfacilitation at the middle knee position. First, at this position, contractile and voluntary knee-extension torque production was at its greatest, presumably placing the highest load on the knee-joint structures, thus stimulating the mechano-sensitive receptors and upregulating afferent feedback by the greatest amount. Indeed, during maximal-effort isokinetic knee-extension contractions at 30°/s, the tibiofemoral joint forces, including the patellar tendon force and joint compressive force, were shown to change with joint angle in proportion with the absolute extension torque production, with the anteriorly-directed shear force (causing anterior tibial translation and thus ACL strain) being the highest at 30-60° of knee flexion (2). These mechanical stimuli would have upregulated signalling via joint (e.g., articular capsule, menisci), ligament, and Golgi tendon organ afferents, which are known to synapse to both Ib inhibitory interneurons (55) and γ-motoneurons (56–58). Together, these effects could result in homonymous Ib inhibition of α-motoneurons, and their disfacilitation due to impaired regulation of muscle spindle sensitivity, leading to attenuation of the excitatory Ia input to the vastii motoneuron pool. Additionally, recurrent inhibition may be stronger at the middle knee position, acting as a safety mechanism preventing further increases in motoneuronal output and thus potentially limiting motor output. This mechanism has been proposed (59) to be responsible for muscle activation failure during maximal eccentric contractions (60, 61), where the intrinsic contractile capacity for force production exceeds isometric capabilities. Finally, reciprocal inhibition could potentially be higher at the middle angle; however, hamstring co-activation has typically been found to be relatively low during isometric knee-extension contractions at both mid-range and extended *vs* flexed knee positions (20, 21, 24).

Furthermore, the augmented upregulation of MU discharge rate required to increase torque output from 50 to 70% MVT at the most flexed knee-joint position (i.e. >optimum length) and consequently higher MU discharge rates during high-level submaximal and maximal voluntary contractions than at the two more extended positions (shorter muscle lengths) suggested a greater degree of motoneuronal discharge facilitation at higher torque levels when the knee is flexed than when it is more extended. Our findings agree with previous observations in soleus and medial gastrocnemius (12), although studies have also reported no effect of joint angle on MU discharge rate modulation with contraction level in tibialis anterior (6, 7) and triceps brachii (8), whereas in hamstrings higher discharge rates at moderate and high torque levels (≥50% MVT) have been shown at short *vs* long muscle lengths (10). These discrepancies may be attributed to relatively small MU samples studied with intramuscular EMG (6, 8), only low contraction levels employed (10 and 20% MVT; (7)), potential differences in neural control between flexors and extensors (62), or possibly to muscle or joint specificity in muscle neural control adjustments with joint angle. Notably, however, our findings are supported by previous observations on the whole-quadriceps level, whereby higher normalised sEMG activity during submaximal isometric contraction levels (23, 24) and its rate of increase with contraction level (23) have been reported to be higher at flexed (long muscle length) compared to extended (short muscle length) knee-joint positions.

The increased motoneuronal output to the vastii muscles at high knee-extension contraction levels and the greater increase in discharge rate with contraction level when the knee is flexed may be attributed to facilitation of the excitatory Ia afferent signalling from the lengthened muscle spindles, which has previously been proposed to explain the greater neuromuscular activation at flexed knee positions (12, 19, 20, 24). Additionally, the inhibitory afferent feedback from knee ligament afferents to the vastii motoneuron pool at 85° of knee flexion is presumably lower due to the largely unloaded ACL and only minimally loaded PCL (3). Whilst this explanation may appear inconsistent with potentially greater levels of reciprocal inhibition of quadriceps from hamstring co-activity, which has generally been found to increase at deep knee-flexion angles (>80°; (20, 21, 24)), it should be noted that the magnitude of reciprocal inhibition reaching the agonist motor pool is modulated by other neural pathways (e.g., descending supraspinal tracts and Renshaw cells) in order to optimise agonist-antagonist coordination according to task demands. The effect of the upregulated hamstring activity in the flexed knee position on quadriceps neural control may therefore not be straightforward and should be investigated with an appropriately designed experiments that could conceivably decouple these mechanisms.

### Intrinsic excitability of spinal motoneurons as a function of knee-joint angle and contraction level

Differential modulation of vastii MU discharge rate with contraction level at the flexed (longest muscle) *vs* both mid-range and extended (shorter muscle lengths) knee positions was also reflected in the estimates of intrinsic excitability of motoneurons. Specifically, increasing the level of triangular contractions from 30 and 50% MVT resulted in both a greater increase in the onset-offset MU discharge rate hysteresis and a smaller decline in the motoneuron input-output nonlinearity at the flexed compared to the two more extended knee positions. This was reinforced by a greater contraction-level-induced gain in the onset discharge rates observed at the flexed *vs* the two more extended positions. Together, these findings suggest a more pronounced upregulation in PIC contribution to motoneuron firing with increased contraction level when the muscle is long compared to when it is at/near the optimal length for torque production or shorter, which may have facilitated the larger MU discharge rate upregulation at high contraction levels. Though speculative, the greater upregulation in spinal excitability may be attributed to a smaller relative increase in local synaptic inhibition with increasing muscle excitation and/or an increased monoaminergic drive from the brainstem.

In addition to the angle-dependent modulation of PICs across contraction levels, we noted a decrease in the onset-offset MU discharge rate hysteresis with increasing degree of knee flexion (increasing quadriceps length), which was consistent with previous observations in human tibialis anterior and gastrocnemius medialis muscles (4) and suggested an overall greater contribution of PIC to motoneuron self-sustained firing when the muscle was short compared to when it was long. In the ankle flexor and extensor muscles this has been attributed to greater levels of spinal reflexive excitability, as previously reported at short *vs* long muscle lengths (63, 64), as well as to possibly lower net inhibitory feedback via joint, ligament, and Golgi tendon organ afferents due to lower absolute torque production and thus internal joint loading at short muscle lengths (4).

### Limitations and further considerations

Some key limitations of this study should be acknowledged. First, MUs were not tracked across contraction levels and/or the knee-joint angles, meaning that samples of MUs studied across different conditions (knee-joint angle and contraction level) likely differed, both in MU number (Table 1) and characteristics (e.g., recruitment threshold). This between-sample variability may have reduced the power of the statistical model to reliably account for the effect of recruitment threshold on MU discharge properties. It is important to recognise, however, that with current decomposition algorithms, MU tracking across a wide range of contraction levels (10-70% MVT) and knee-joint angles would likely be limited due to changes in muscle geometry and thus MU action potential shape as a function of both contraction level (65) and muscle length (66). Second, during MVCs the recruitment threshold of active MUs cannot be reliably determined due to the high rate of torque development and thus MU recruitment threshold could not be included in the statistical model as a covariate in this specific analysis, which may have underestimated the estimated marginal means of MU discharge rate during MVCs. This is particularly problematic in cases with low MU yields, typically comprising high-threshold MUs (due to the decomposition bias) with relatively lower MU discharge rates (according to the onion skin principle). This limitation could be overcome by tracking MUs from lower-level ramp contractions (where the recruitment threshold can be more reliably determined) to MVCs. However, this approach has its own limitations, including biasing the MU sample to MUs with recruitment thresholds within the torque range of the ‘calibration’ contraction(s) and potentially limiting the capacity to identify the lower-threshold MUs during MVCs due to MU superimposition. Finally, it should be noted that in this study, only a limited range (i.e., 60°) of the entire functional range of motion of a healthy knee (∼150°) has been studied. Whether vastii MU discharge rates and contraction-level induced gain in motoneuron excitability continue to increase, plateau, or possibly decrease at more flexed angles (very long quadriceps lengths) remains to be explored.

### Conclusion

In conclusion, we showed that neural control of vastii muscle torque production during submaximal and maximal isometric knee-extension contractions varies with the knee-joint angle (muscle length). Specifically, MU discharge rates were the lowest at the intermediate knee joint position, where contractile and voluntary torque production capacities were the greatest, which is likely due to the combination of high absolute muscle torque and internal joint loading leading to greater inhibitory input to the spinal cord that limits motoneuronal output. In contrast, MU discharge rates during high-level submaximal and maximal contractions were the highest at the most flexed knee position, stemming from a greater upregulation of motoneuronal output with increasing torque at higher contraction levels, likely due to a greater contraction-level-induced increase in PIC magnitude. These findings suggest the balance between excitatory and inhibitory sensory feedback to the central nervous system differs between knee-joint positions and significantly modulates neural command to the extensor muscles. This emphasises the importance of considering knee-joint angle when studying quadriceps neural control in healthy individuals or when examining adjustments to chronic conditions (e.g., knee osteoarthritis) or following acute knee injuries (e.g., ACL injury).

## ACKNOWLEDGEMENTS

This activity was conducted under the auspices of the National Rehabilitation Centre (NRC), a collaboration between Loughborough University, The University of Nottingham and Nottingham University Hospitals NHS Trust. The views expressed are those of the authors and not necessarily those of the NRC or the partners involved.

